# Understanding the B and T cells epitopes of spike protein of severe respiratory syndrome coronavirus-2: A computational way to predict the immunogens

**DOI:** 10.1101/2020.04.08.013516

**Authors:** Yoya Vashi, Vipin Jagrit, Sachin Kumar

## Abstract

The 2019 novel severe respiratory syndrome coronavirus-2 (SARS-CoV-2) outbreak has caused a large number of deaths with thousands of confirmed cases worldwide. The present study followed computational approaches to identify B- and T-cell epitopes for spike glycoprotein of SARS-CoV-2 by its interactions with the human leukocyte antigen alleles. We identified twenty-four peptide stretches on the SARS-CoV-2 spike protein that are well conserved among the reported strains. The S protein structure further validated the presence of predicted peptides on the surface. Out of which twenty are surface exposed and predicted to have reasonable epitope binding efficiency. The work could be useful for understanding the immunodominant regions in the surface protein of SARS-CoV-2 and could potentially help in designing some peptide-based diagnostics.

## Introduction

Emerging severe acute respiratory syndrome coronavirus 2 (SARS-CoV-2) is a recent pandemic and declared as a public health emergency by World Health Organization (WHO) ^1^. The disease rapidly spread across the globe and caused havoc to humanity ^2^. By the end of March, SARS-CoV-2 had spread to 200 countries and infected over 4,50,000 people ^3^. The WHO is continuously monitoring and updating the health-related plans to curtail the disease spread. The absence of specific treatment and vaccine worsen the situation and threat the world.

International Committee on Taxonomy of Viruses (ICTV), classified SARS-CoV-2 under family *coronaviridae* of order *nidovirales*. The genomic sequence of SARS-CoV-2 isolated from the bronchoalveolar lavage fluid of a patient from the Wuhan, China showed a length of 29,903 nucleotides (GenBank accession number NC_045512) ^4^. The SARS-CoV-2 contains a positive single-stranded RNA with 5’ and 3’ UTR. The genome codes for ORF1a, ORF1b, Spike (S), ORF3a, ORF3b, Envelope (E), Membrane (M), ORF6, ORF7a, ORF7b, ORF8, ORF9b, ORF14, Nucleocapsid (N), and ORF10 from 5’ to 3’ ^4,5^.

The S glycoprotein forms homotrimers and represents a potential target for therapeutic and vaccine design as it mediates viral entry into host cells ^6,7^. S glycoprotein comprises of two functional subunits. Whereas the S1 subunit is responsible for binding to the host cell receptor, the S2 subunit is responsible for the fusion of the viral with the cell membrane. Usually, in CoVs, S is cleaved at the boundary between the S1 and S2 subunits, which remain non-covalently bound in the prefusion conformation, to activate the protein for membrane fusion via extensive irreversible conformational changes ^8-10^. Setting apart from other SARS-CoVs, it is found that S glycoprotein of SARS-CoV-2 harbors a furin cleavage site at the boundary between the S1/S2 subunits ^11^. By now, it is evident that SARS-CoV-2 S uses angiotensin-converting enzyme 2 (ACE2) receptor-mediated entry into cells. It is found that the receptor-binding domains of S proteins of SARS-CoV-2 and SARS-CoV bind with similar affinities to human ACE2 ^11,12^.

As the situation worsens, there is a growing need for the development of suitable therapeutics and alternate diagnostics against SARS-CoV-2 for effective disease management strategies. Diagnostic assays based on peptides have become increasingly substantial and indispensable for its advantages over conventional methods ^13^. The present study aimed to locate appropriate epitopes within a particular protein antigen, which can elicit an immune response that could be selected for the synthesis of the immunogenic peptide. Using computational approach, S glycoprotein of SARS-CoV-2 was explored to identify various immunodominant epitopes for the development of diagnostics. Besides, the results could also help us to understand the SARS-CoV-2 surface protein response towards T and B cells.

## Materials & Methods

### Collection of targeted protein sequence

We downloaded amino acid sequences (n=98) of S protein available at the time of study on targeted SARS-CoV-2 from the National Centre for Biotechnological Information (NCBI) database.

### Identification of potential peptides

To identify an immunodominant region, it is of extreme importance to select the conserved region within the S protein of SARS-CoV-2. All the sequences were compared among themselves for variability using protein variability server by Shannon method ^14^. The average solvent accessibility (ASA) profile was predicted for each sequence using SABLE server ^15^. BepiPred 1.0 Linear Epitope Prediction module ^16-18^ incorporated in Immune Epitope Database (IEDB) ^19^ was used to predict potential epitopes within the S protein. The FASTA sequence of the targeted protein was used as an input for all the default parameters.

### Identification of B-cell epitopes

We used two web-based tools for B-cell epitope prediction, viz., the IEDB, and ABCpred server ^20^. S protein structure from protein data bank (PDB) (6VSB) ^21^ was analyzed for linear and discontinuous B-cell epitopes using ElliPro module ^22^ on IEDB server, with default settings. Also, ABCpred server was used to detect B-cell epitopes using artificial neural network (ann) method.

### Identification of T-cell epitopes

T-cell epitopes having binding affinity towards MHC-I and MHC-II alleles were selected to boost up both cytotoxic T-cell and helper T-cell mediated immune response. IEDB server was used to predict the MHC-I and MHC-II binding epitopes for targeted protein. The reference set of alleles was used for predicting the MHC-I and MHC-II T-cell epitopes. ^23-27^

## Results and Discussion

In our study, we targeted the S glycoprotein of SARS-CoV-2 as it is present outside the virus and interacts with the host receptor. At the time of the study, there were 98 sequences available for the targeted protein of SARS-CoV-2. The S glycoprotein is 1273 amino acids long sequence except for the virus isolated from Kerala (India), which is 1272 amino acids long spike glycoprotein (GenBank accession number MT012098). Our interest here was to determine conserved regions first and then determine surface-exposed regions, which are potential epitopes to generate immune response. We found that sequences among all the S proteins in the analysis are least variable and highly conserved as shown in Figure 1. Regions having a high value of ASA are more surface exposed as compared to others. We identified a total of 24 peptides of varying length, as shown in Table 1, which are selected based on high ASA values. The potential epitope regions were predicted using the sequence of S protein of SARS-CoV-2, which showed the least variability (GenBank accession number NC_045512). The potential epitopes are represented by blue peaks, while green-colored slopes represent non-epitopic regions (Figure 2).

**Table 1.**
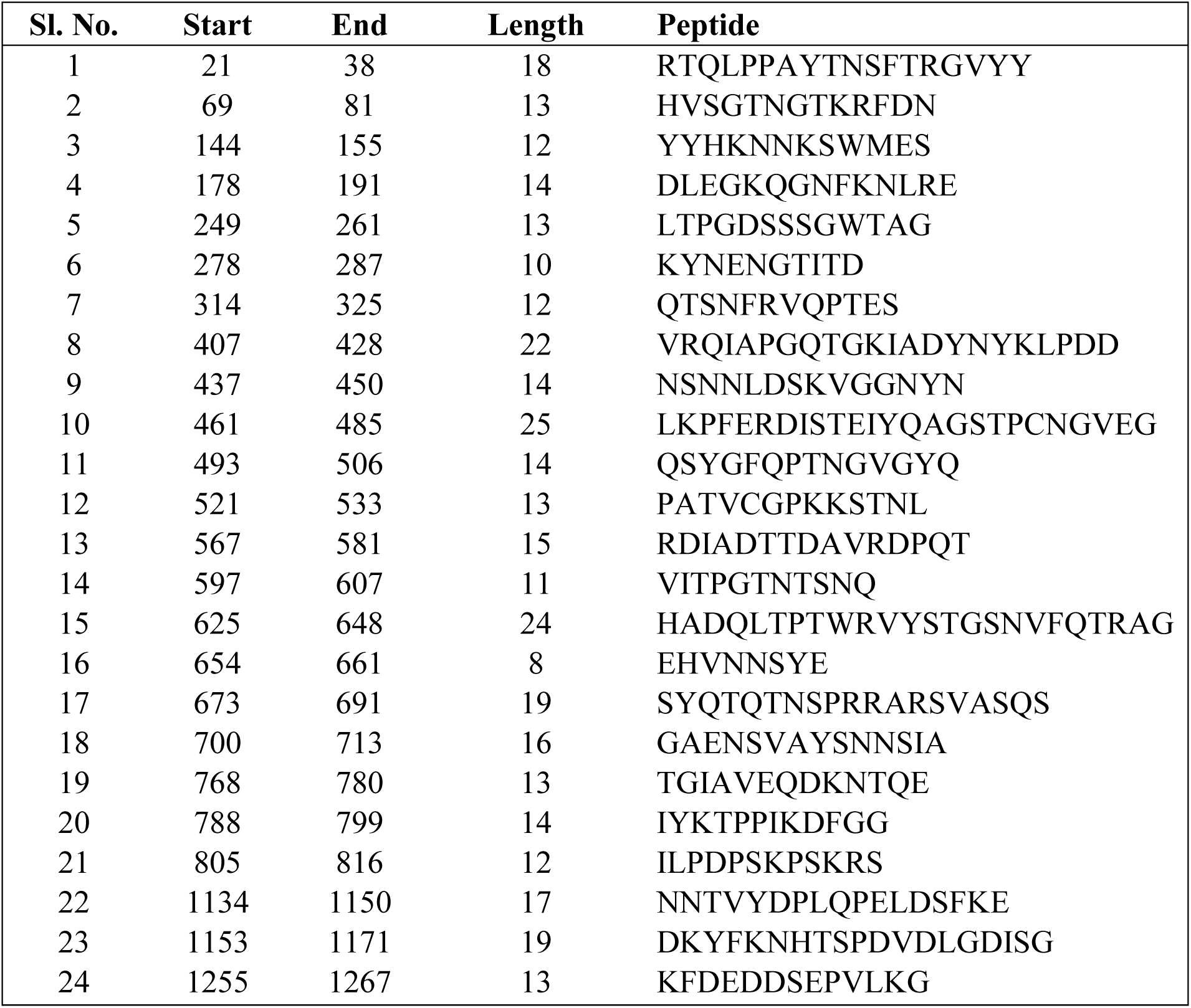
Conserved region with good average solvent accessibility selected for further analysis.

**Figure 1.**
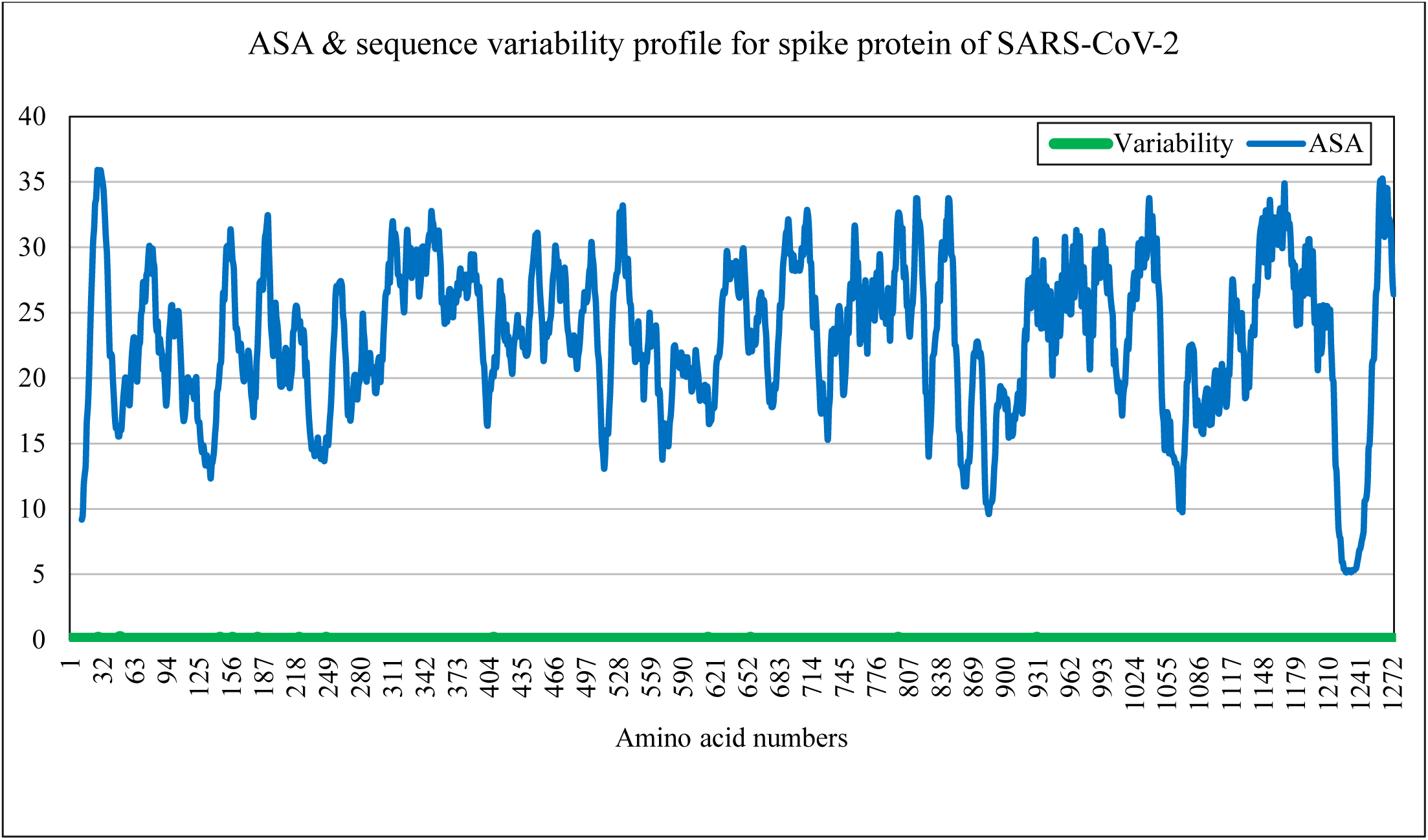
Profiles of average solvent accessibility (blue) in % and amino acid sequence variability (green) in numbers of the 98 SARS-CoV-2 protein plotted against amino acid numbers.

**Figure 2.**
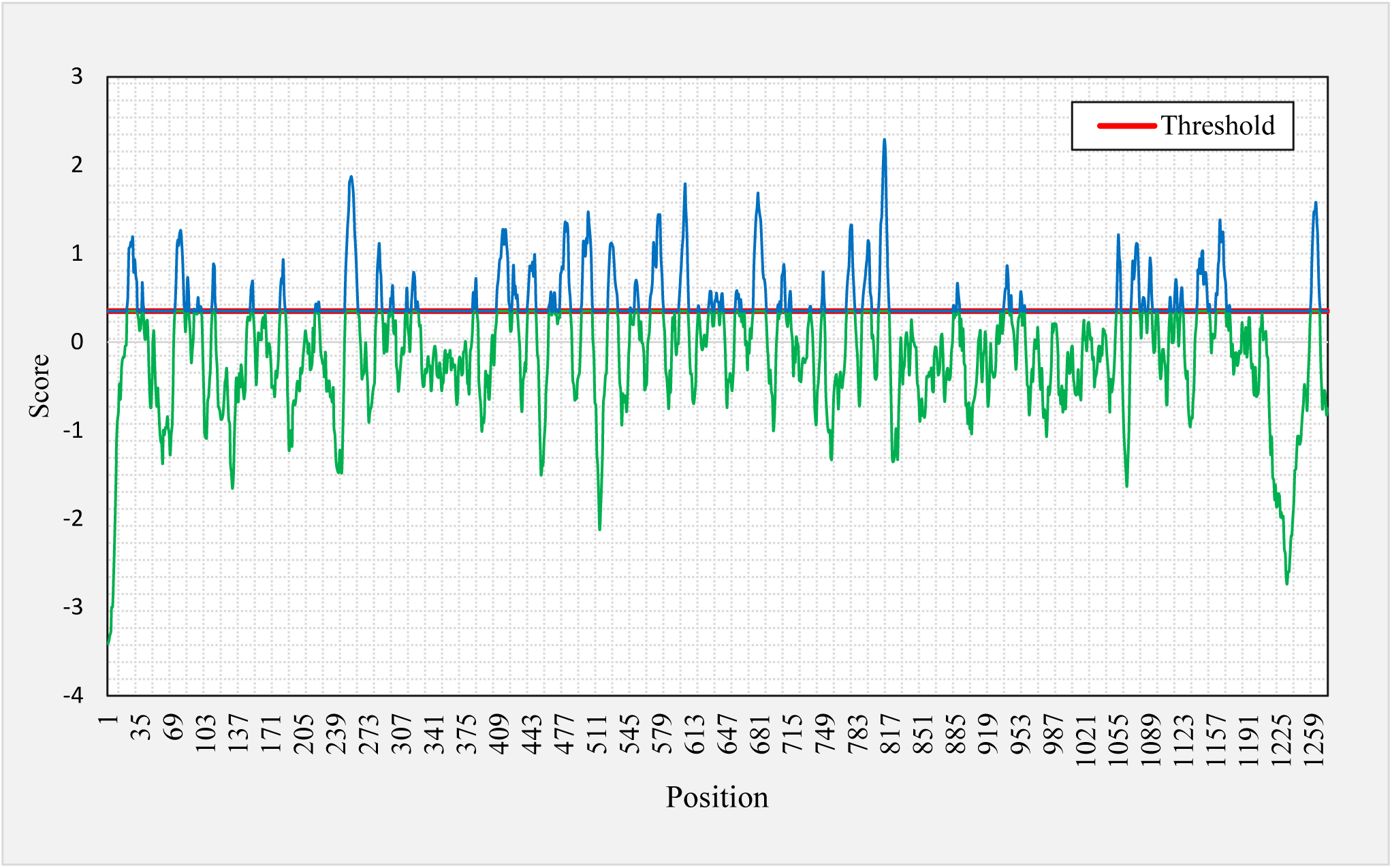
Graphical representation of B-cell linear epitopes of spike protein of SARS-CoV-2. B-cell linear epitopes predicted using BepiPred 1.0 module incorporated in IEDB server using default threshold value (0.35).

The existence of B-cell linear and discontinuous (conformational) epitopes within the identified segments could help us to identify the peptides, which can elicit immune response ^28^. We identified 18 linear epitopes, predicted by ElliPro (IEDB), which contains regions from 19 of our selected peptides highlighted in red in Table 2. These identified B-cell linear epitopes are placed based on their positional value, and scores. Epitopes with high scores have more potential for antibody binding. Five of our selected peptides (peptide numbers 3, 5, 19, 23, and 24 in Table 1) were not considered as potential linear B-cell epitopes.

**Table 2.**
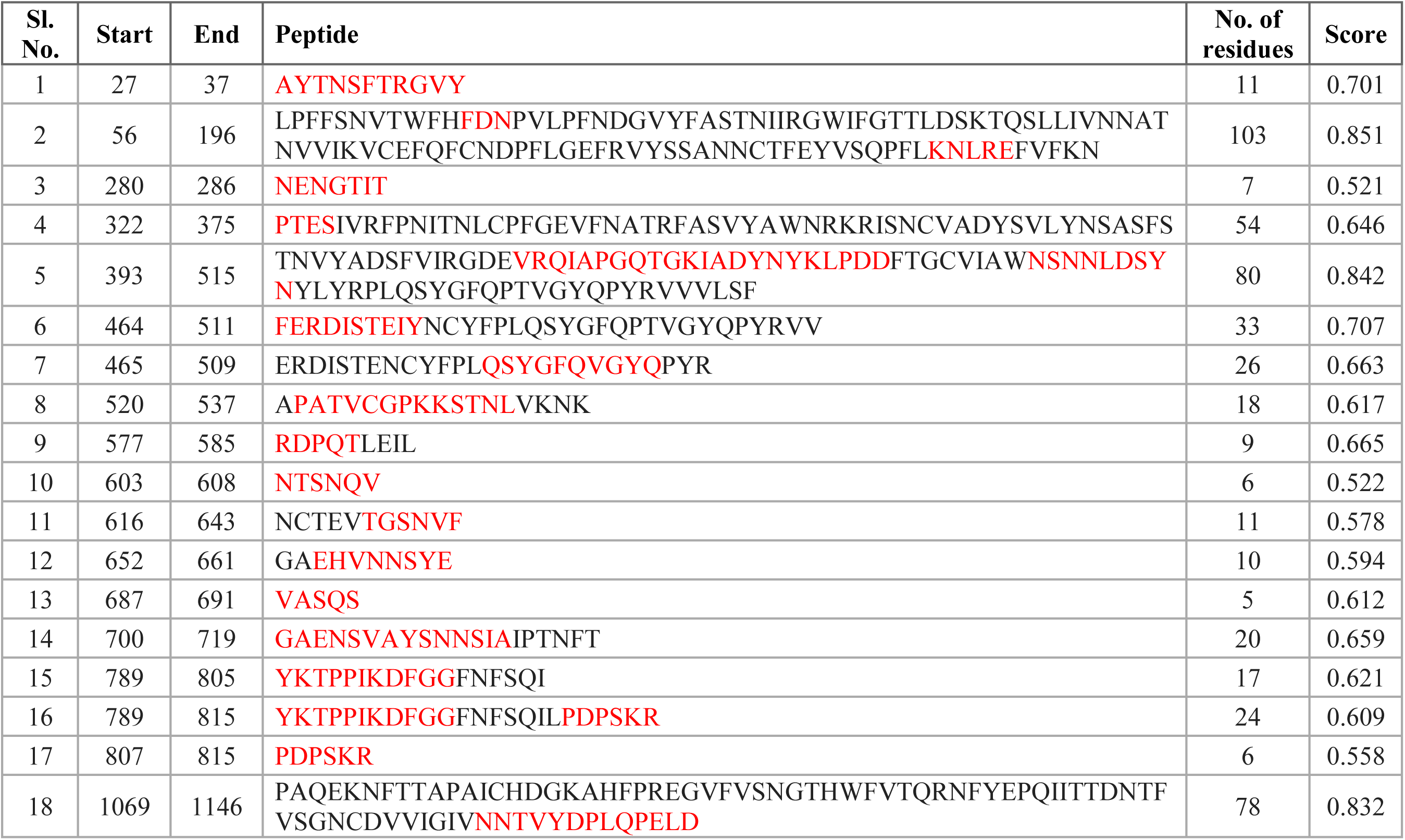
IEDB ElliPro predicted linear epitopes for spike protein of SARS-CoV-2. Sequences that match our selected peptides are marked in red.

Using the same module, B-cell discontinuous epitopes were predicted, which gave 16 epitope regions that contained regions from 18 of our selected peptides highlighted in red (Table 3). Six peptides (peptide numbers 3, 5, 14, 19, 23, and 24 in Table 1) were not predicted as discontinuous B-cell epitopes. To further confirm, we used ABCpred server to detect B-cell epitopes, with default threshold of 0.51. It identified various epitopes with different length and scores; out of those, the regions which contained our selected peptides are highlighted in red (Table 4). A high score represents a good binding affinity with epitopes, and most of our peptides scored more than 0.7 and were predicted as linear B-cell epitopes.

**Table 3.**
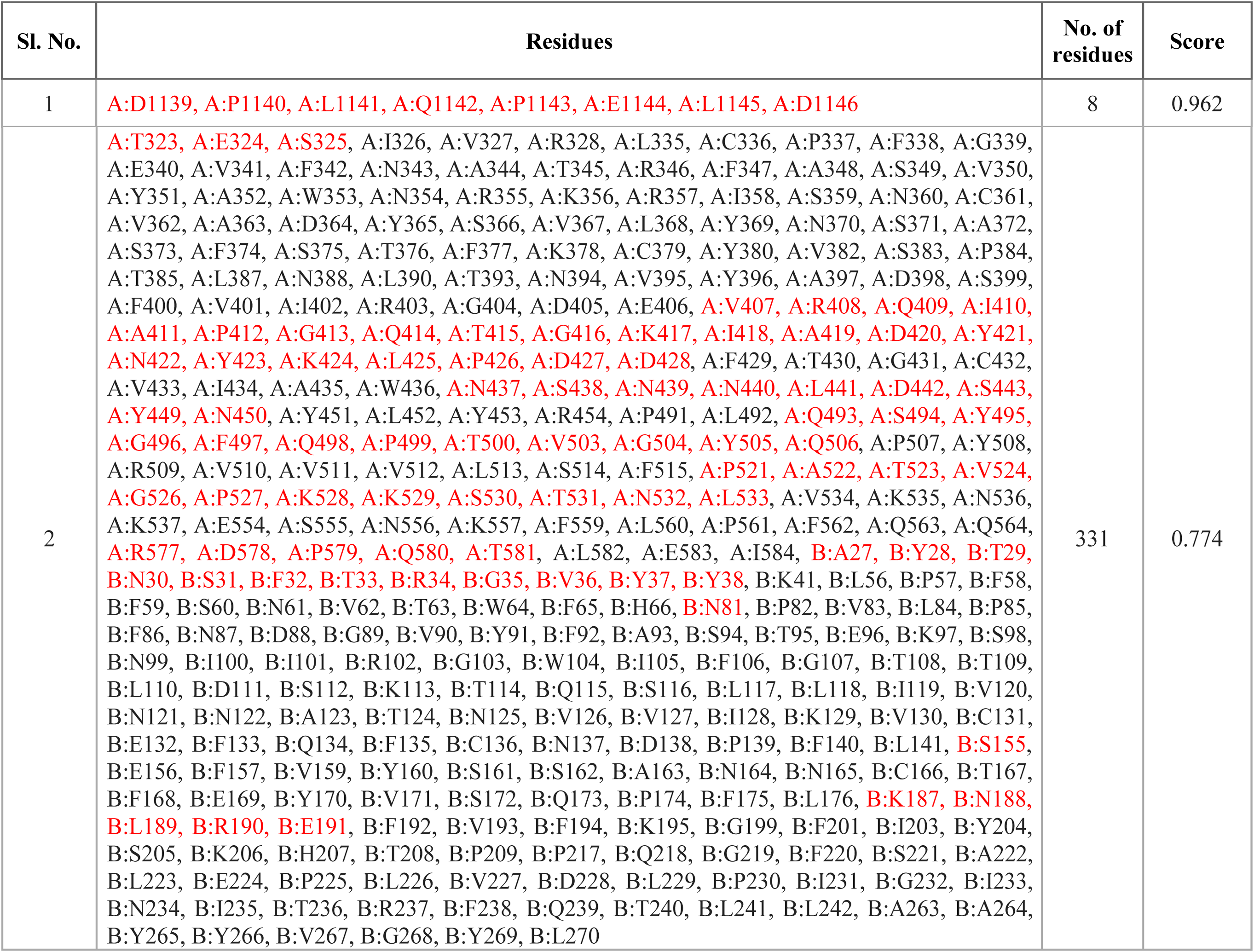
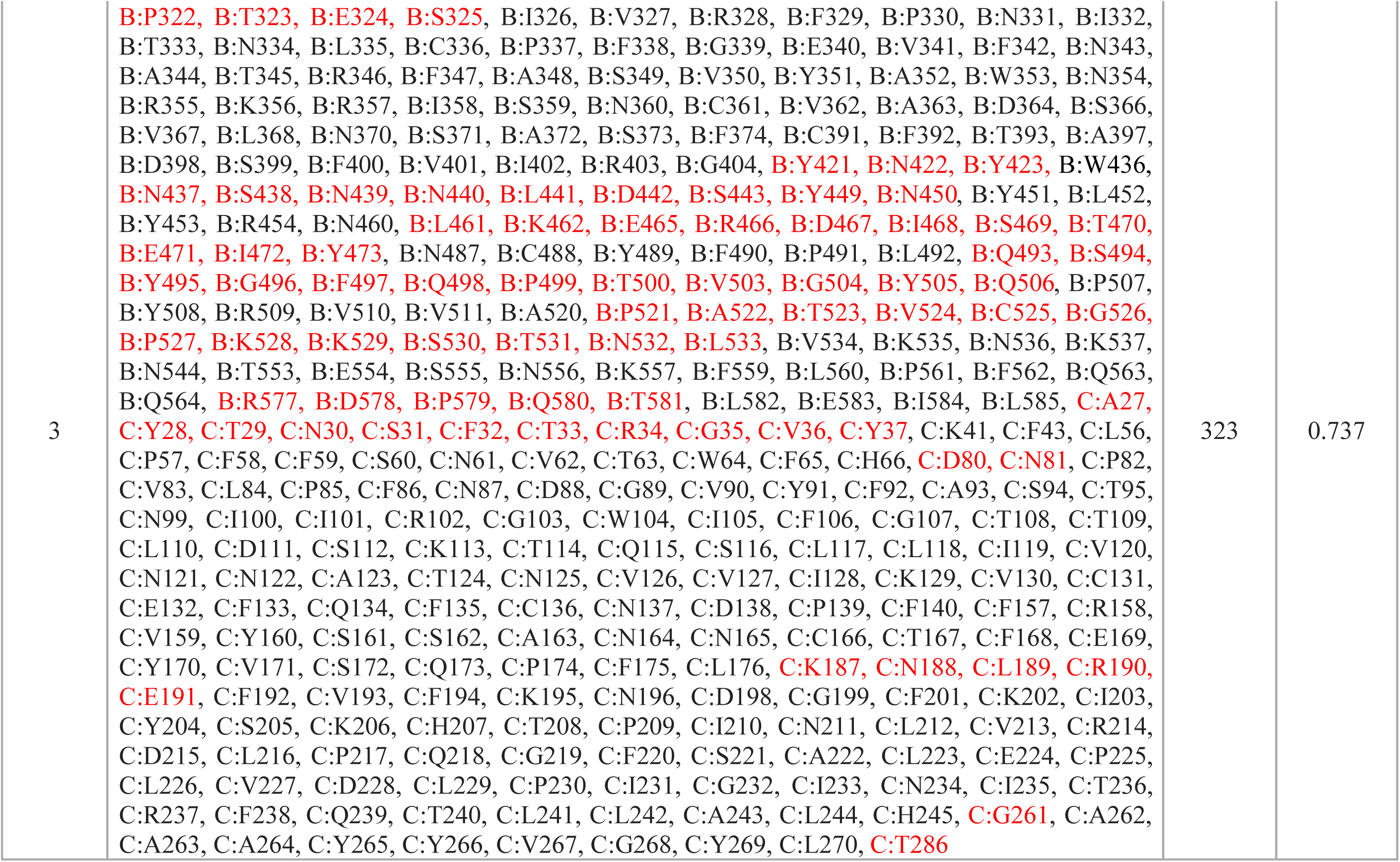
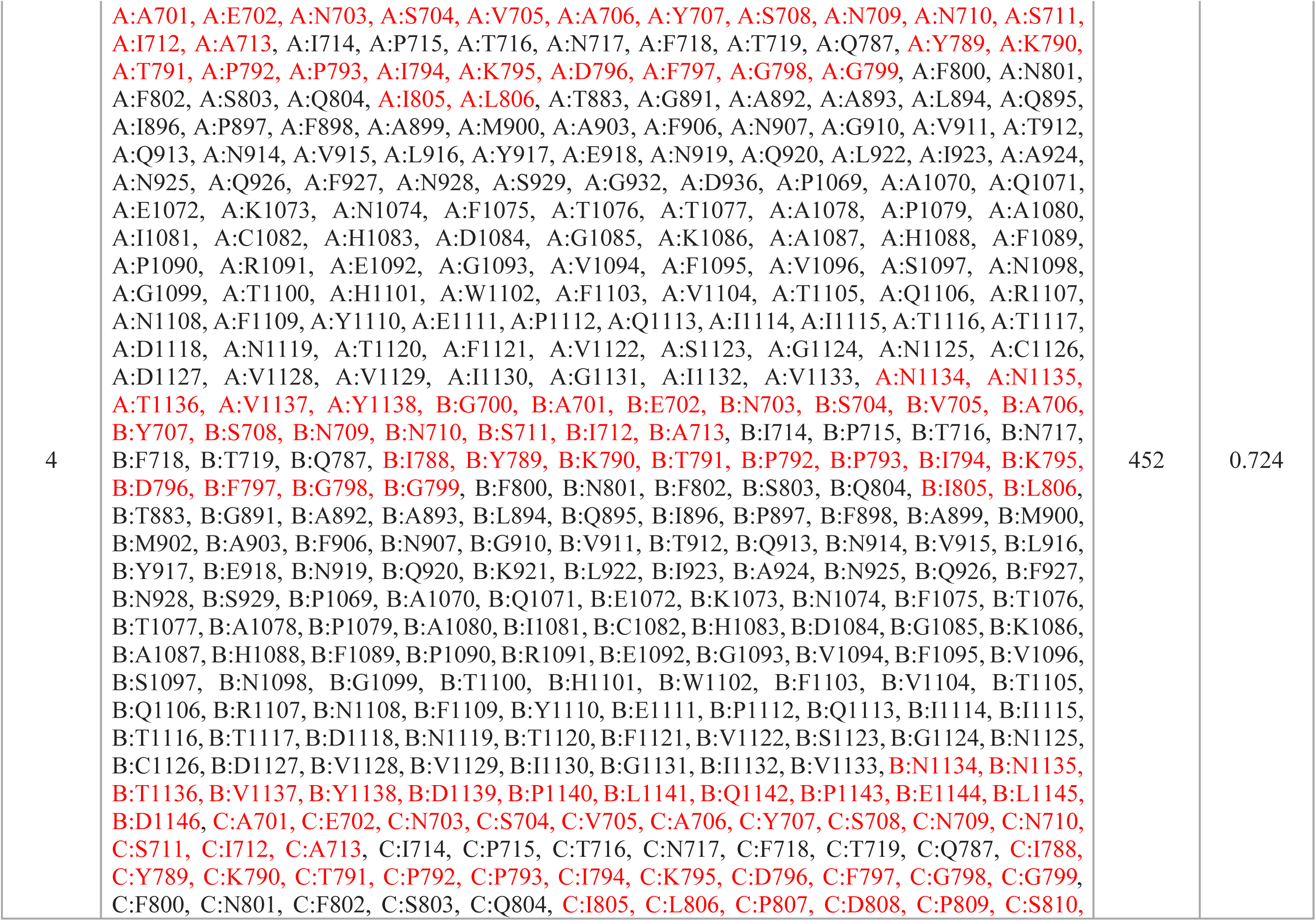
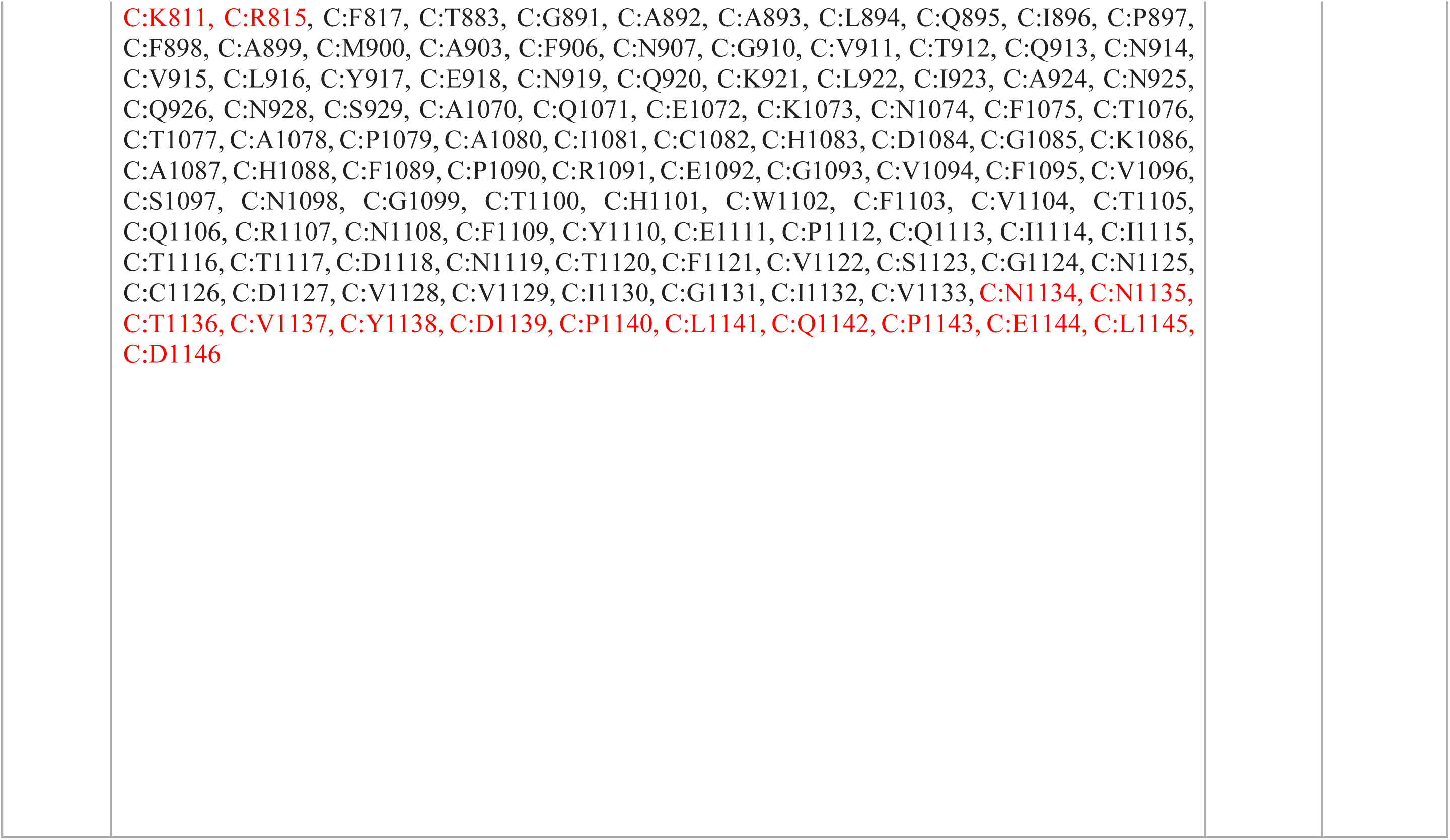
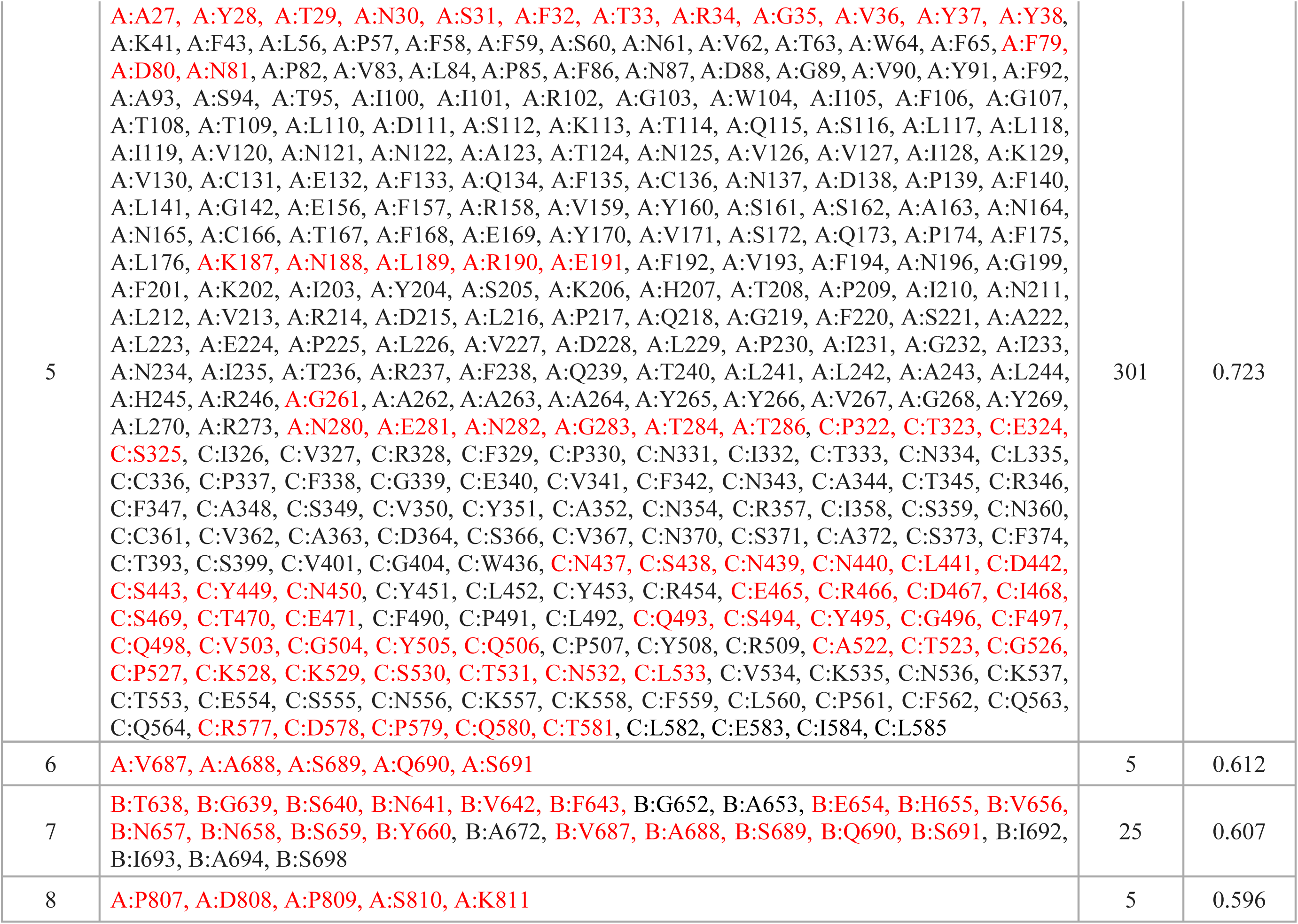
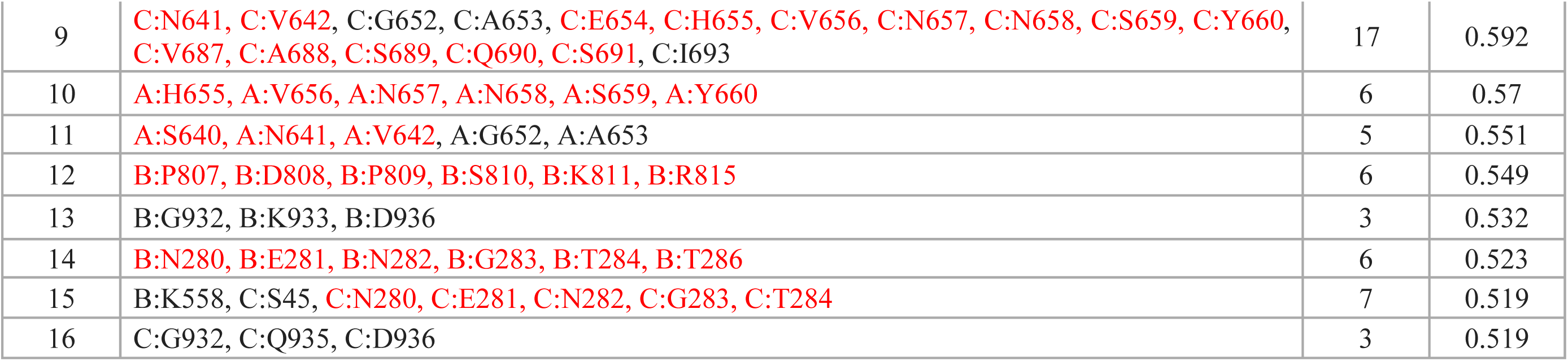
IEDB ElliPro predicted discontinuous epitopes for spike protein of SARS-CoV-2. Sequences that match our selected peptides are marked in red.

**Table 4.**
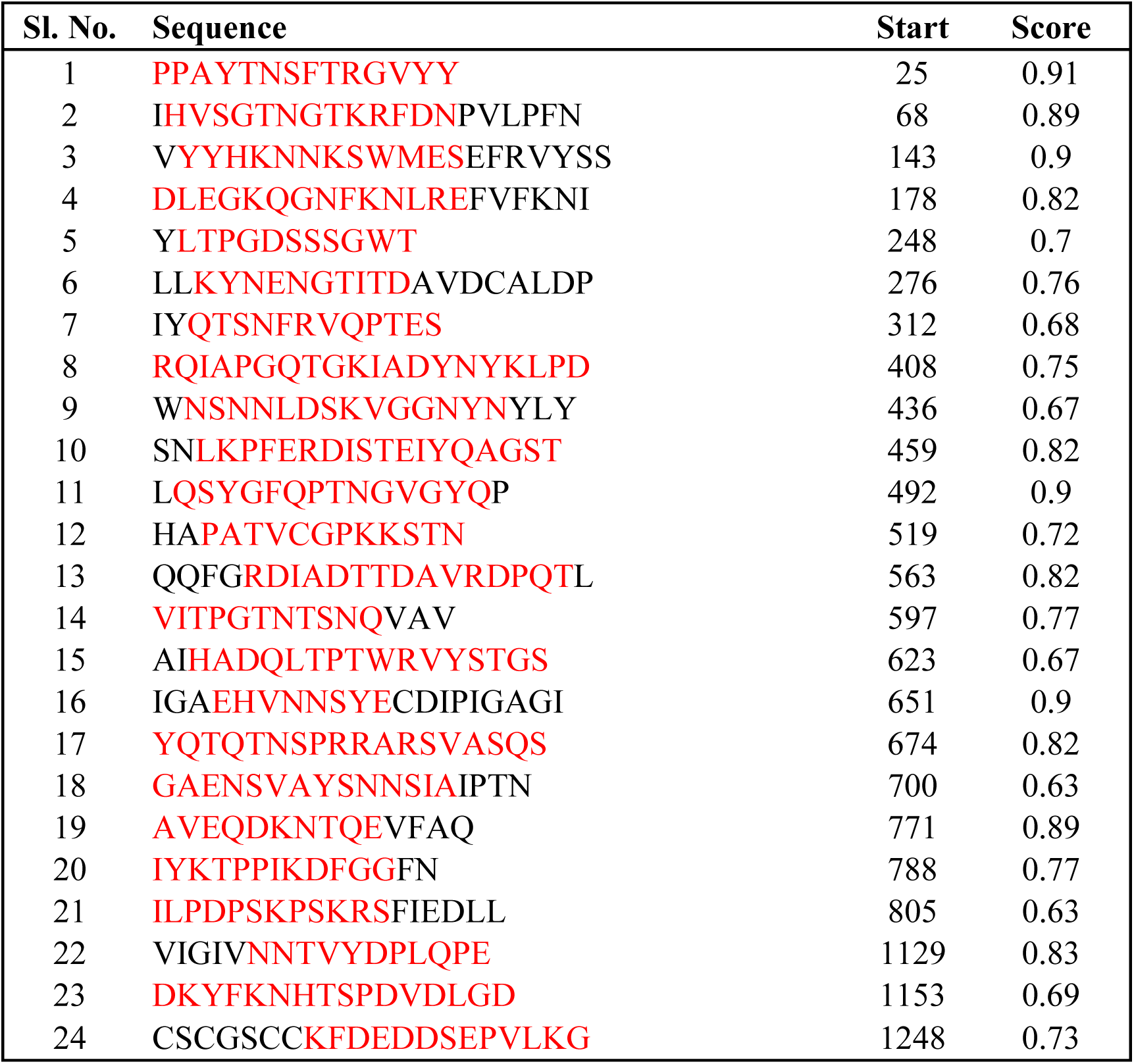
ABCpred determination of B-cell binding affinities. Note that high score indicates good binding affinity.

We used the IEDB server to determine the binding affinity for human leucocyte antigen (HLA) with our selected peptides from Table 1. As recommended by the IEDB server, reference HLA allele sets were used for the prediction of MHC-I and MHC-II T-cell epitopes, as they provide comprehensive coverage of the population. All the predictions were made using IEDB recommended procedures. We observed good binding affinities for our selected peptides. The list of binding affinities for MHC-I T-cell epitopes is given in Table 5, where low rank represents high binding affinity. The epitopes with rank <1% for very high binding affinity were selected. Regions from all of our selected peptides were found to be potential T-cell epitope(s) with high binding affinity with HLA-A and HLA-B alleles, except one. Similarly, the list of binding affinities for MHC-II T-cell epitopes are given in Table 6. Regions from our selected peptides are highlighted in red. The results revealed that around half of our selected peptides are potential T-cell epitope(s) with high binding affinity with HLA-DRB and HLA-DP/DQ alleles.

**Table 5.**
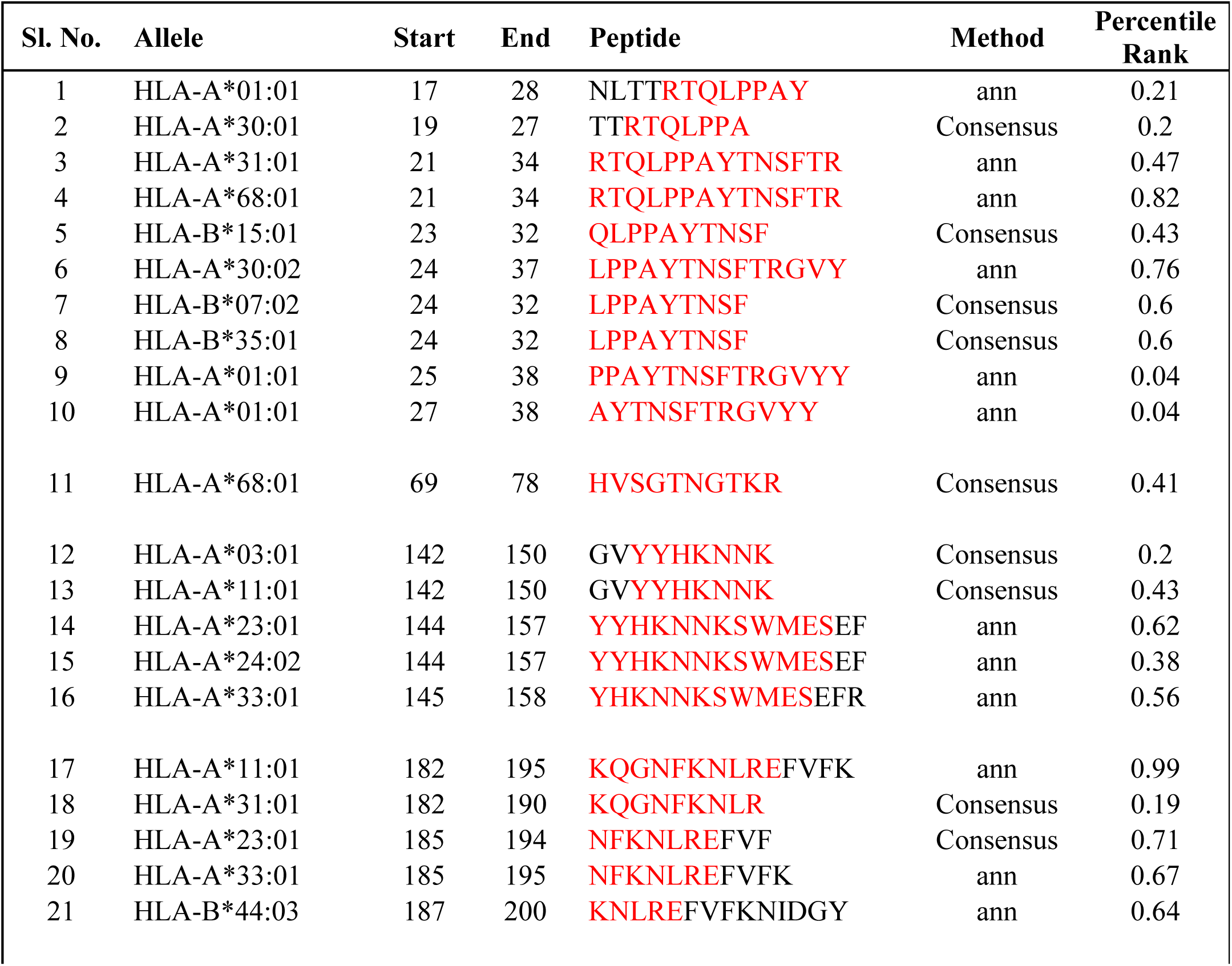
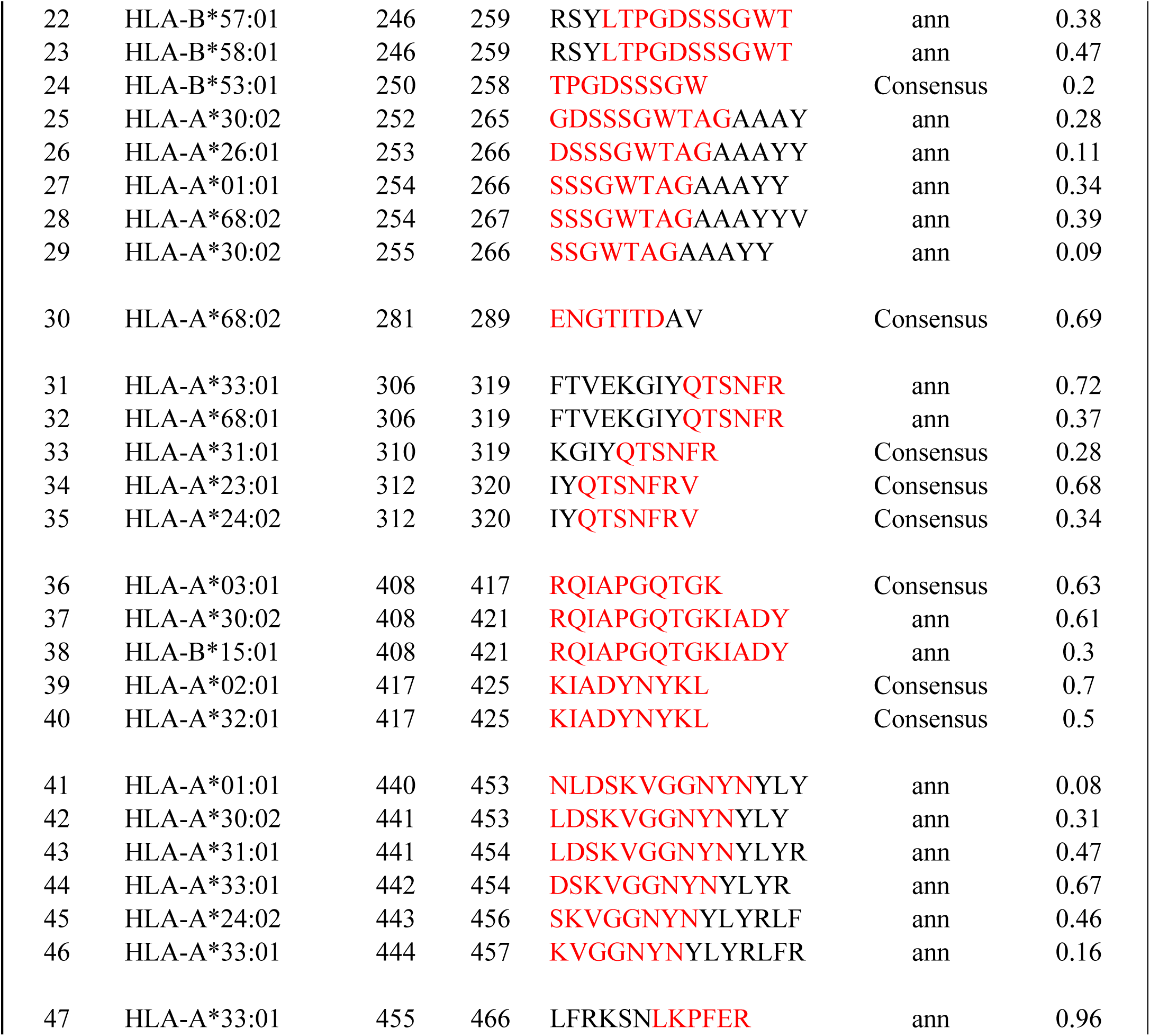
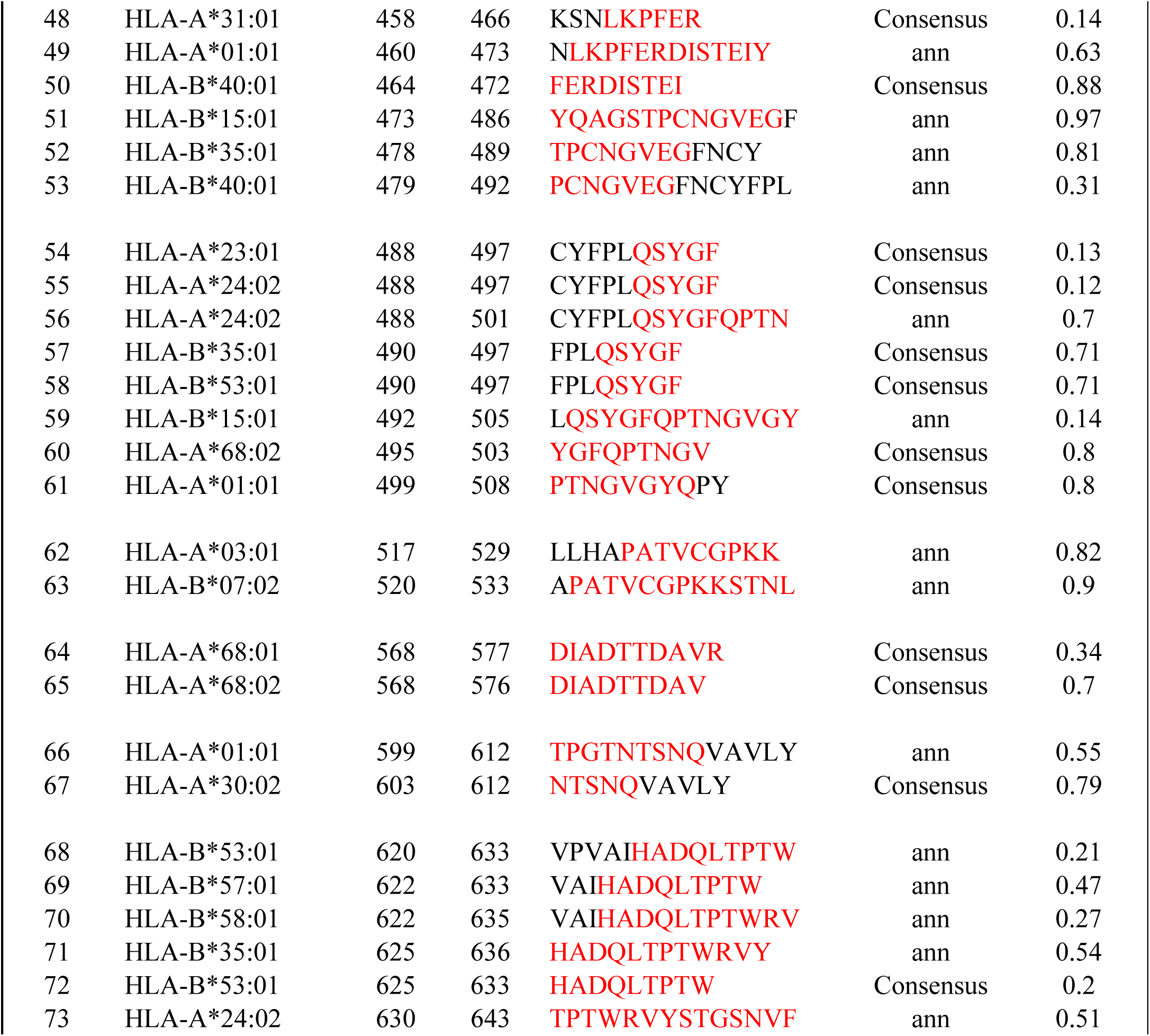
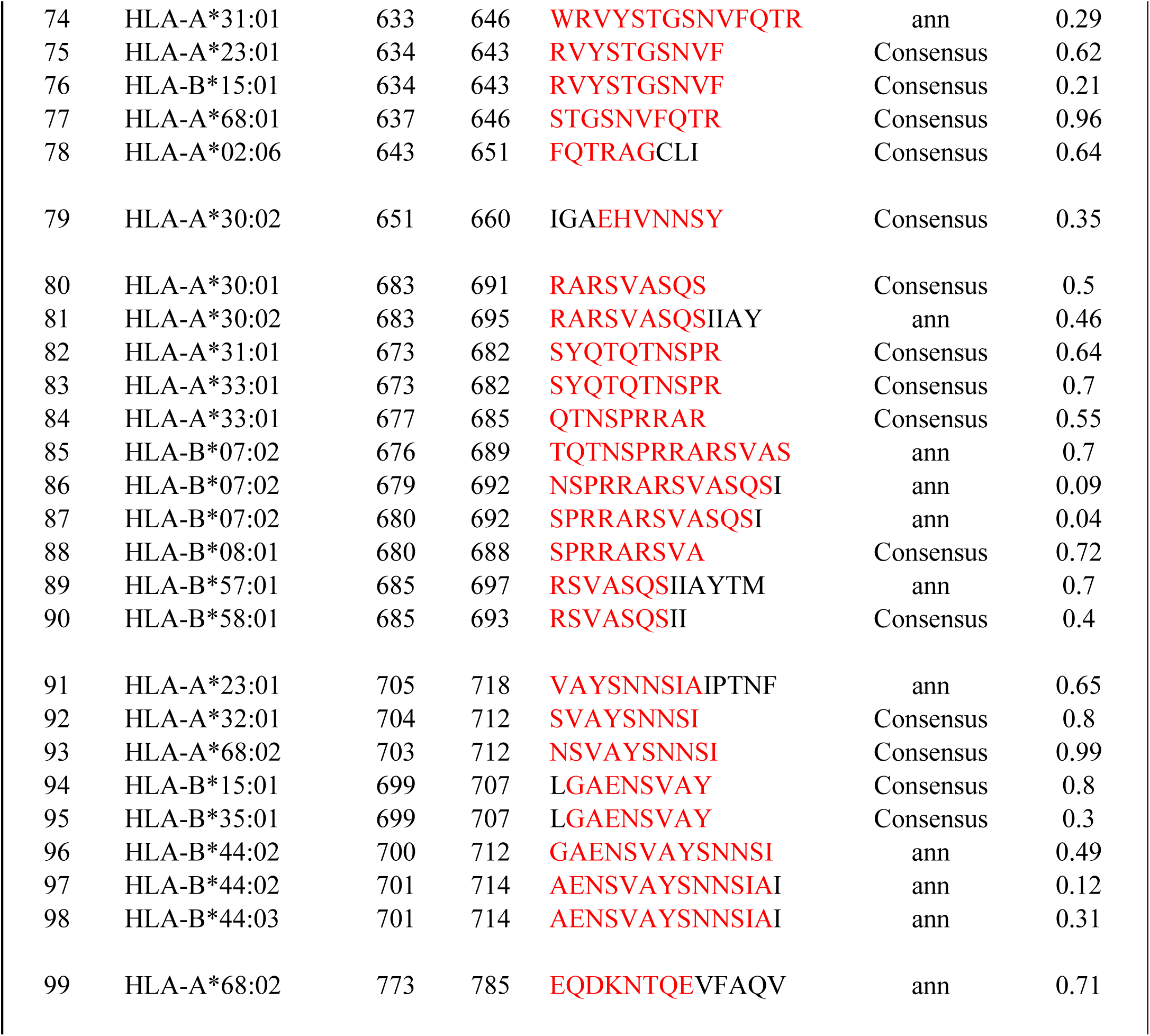
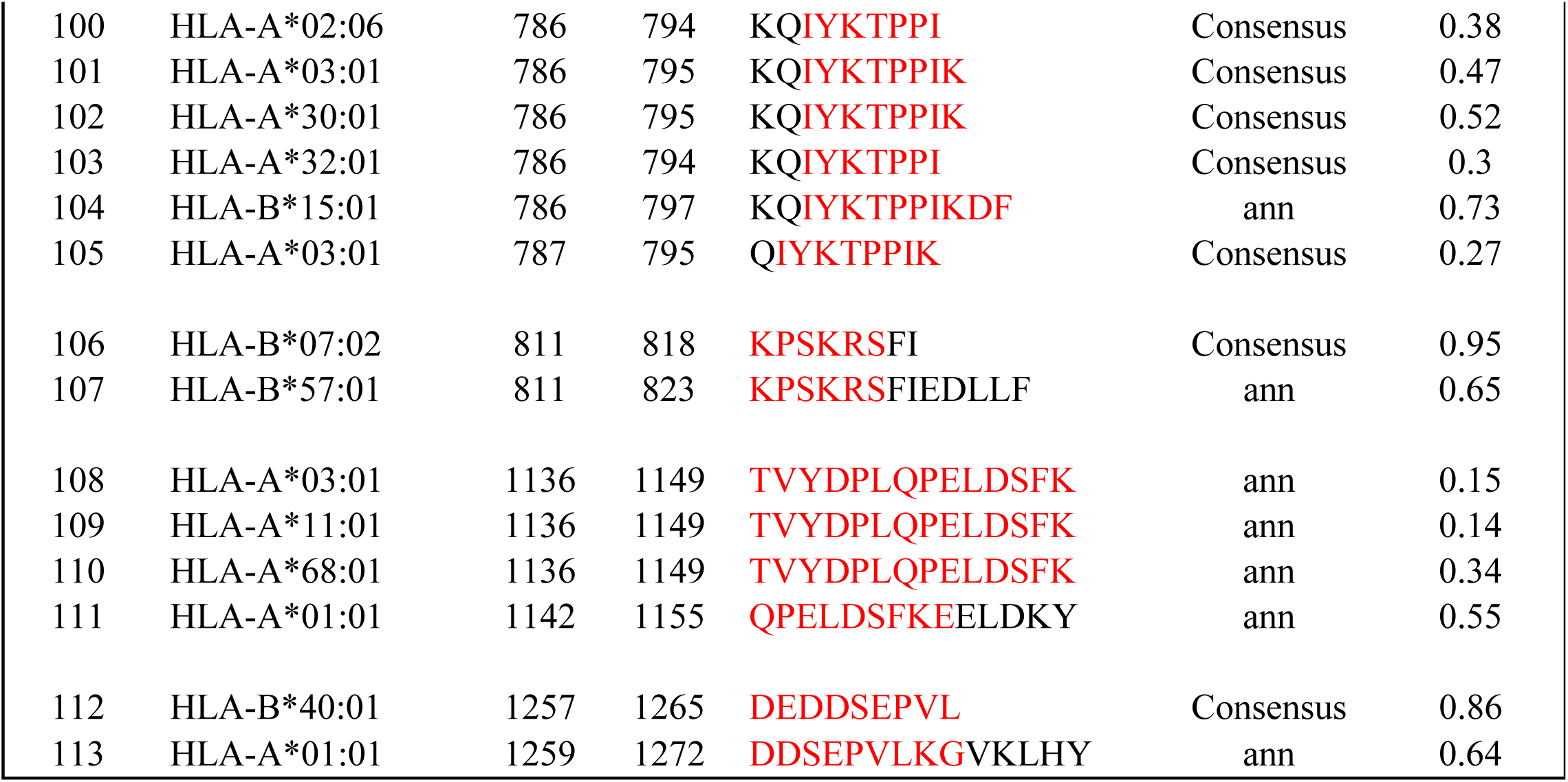
IEDB prediction of binding affinity with MHC-I alleles, only our selected peptides with percentile rank less than 1.00 are shown here. The binding affinity is considered higher for low percentile rank. Sequences that match our selected peptides are marked in red.

**Table 6.**
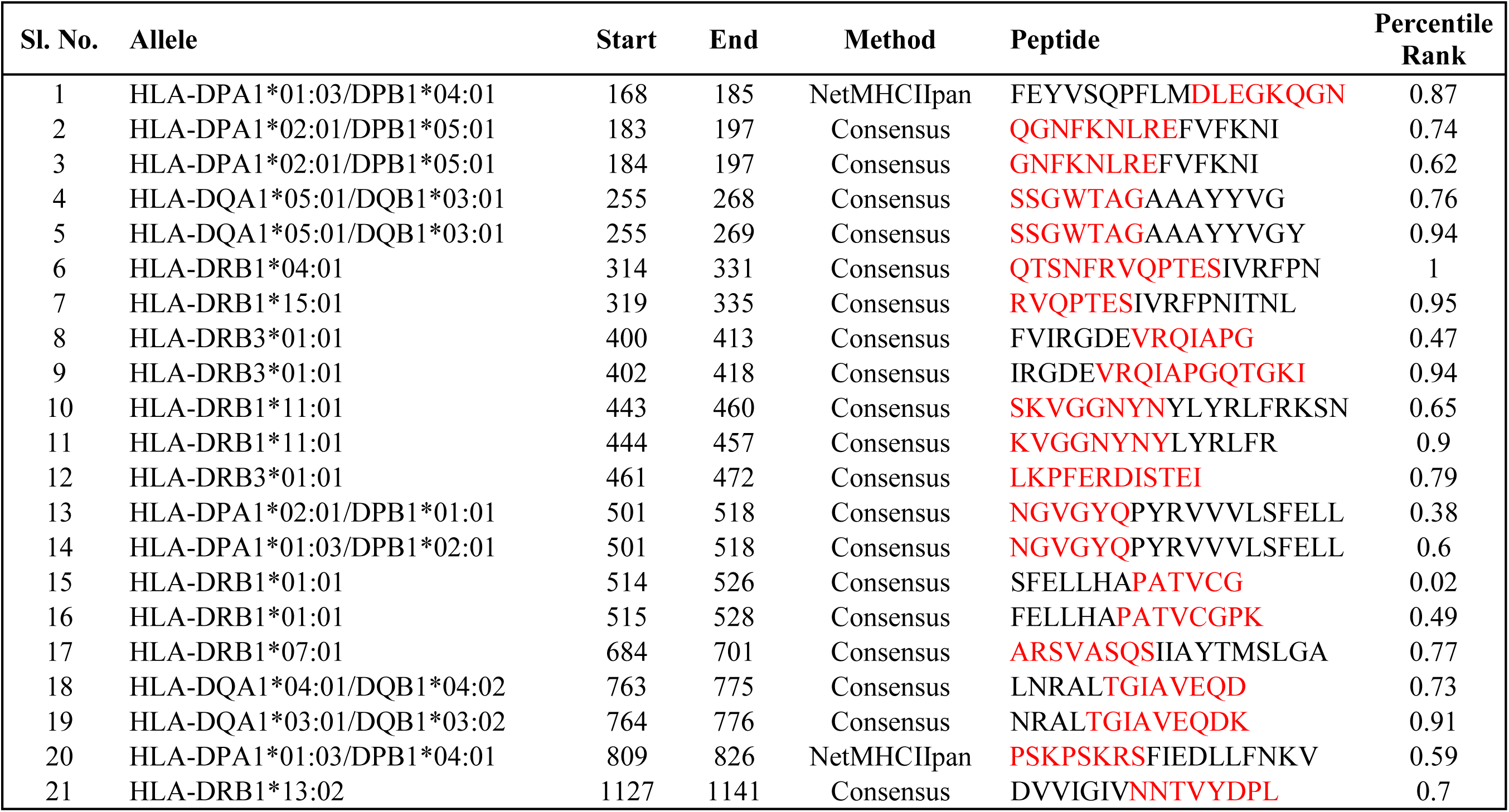
IEDB prediction of binding affinity with MHC-II alleles, only our selected peptides with percentile rank less than 1.00 are shown here. The binding affinity is considered higher for low percentile rank. Sequences that match our selected peptides are marked in red.

Overall, it was found that the regions identified in Table 1 not only had good B-cell and T-cell affinities, but the majority of them had overlaps with discontinuous epitopes also (Table 3). The peptide segments identified from the set of 98 sequences of the SARS-CoV-2 S glycoprotein appear to hold reasonable potential to act as immunogens. Peptide-based diagnostics and vaccines have been proposed against virus outbreaks earlier ^29-33^. The availability of a 3D structure (6VSB) of the SARS-CoV-2 S glycoprotein provided an opportunity to inspect the predicted peptides. Placement of the peptide segments identified by ASA and conserved sequence analysis on the S glycoprotein showed that 20 regions that we identified lie on the surface (Figure 3). In order to limit recognition and evade from immune response of host, coronaviruses use conformational masking and glycan shielding ^34,35^. SARS-CoV-2 S trimer also exists in multiple distinct conformational states, which is necessary for receptor engagement leading to initiation of fusogenic conformational changes ^11^. A considerable good number of peptides at the surface region of the S glycoprotein allows the potential use of those peptide regions as immunogens.

**Figure 3.**
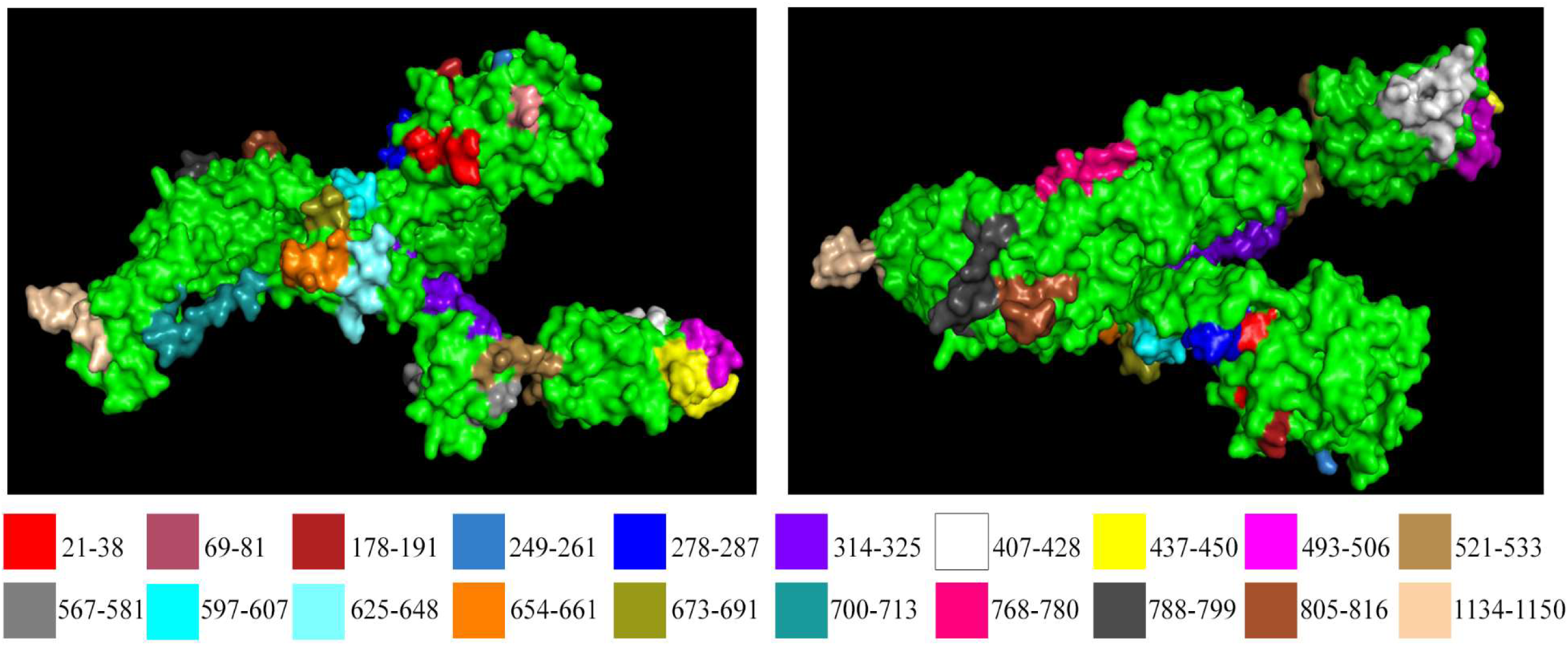
Our selected peptides are highlighted on spike protein of SARS-CoV-2 protein structure downloaded from PDB (ID: 6VSB).

The emergence of new viral diseases like SARS-CoV-2 represents a substantial global disease burden. There is an urgent need for diagnostics, therapeutics, and vaccines against newly emerged SARS-CoV-2. Facilitated by high mutation rates, traditional vaccines based on antibody-mediated protection are often poor inducers of T cell responses and can have limited success ^36^. In our study, we predicted both B-cell and T-cell epitopes for conferring immunity in different ways. We speculate that the identified epitopes with considerably good epitope binding efficiency have the potential to be an immunodominant peptide. Peptide-based sensitive and rapid diagnostic kits are considered as a better alternative to the conventional serological tests including whole antigenic protein ^13^. The study could help us to use the predicted peptide as an immunogen for the development of diagnostics against SARS-CoV-2.

## Conclusion

In the present study, peptide segments were identified on S proteins for the development of diagnostics against SARS-CoV-2. The recent availability of 3D data on 2019-CoV spike glycoprotein has helped the search. SARS-CoV-2, being an RNA virus, has high mutations rate and undergoing active recombination ^37^. Although the peptides identified are ideal candidates as immunogens for peptide-based diagnostics development, more refinement and lab trials are essential steps that are yet to be undertaken for early development of diagnostics before the identified epitopes are rendered obsolete.

